# Presenilin-dependent ErbB4 Nuclear Signaling Regulates Astrogenesis via PARP1

**DOI:** 10.64898/2026.09.22.753567

**Authors:** Mostafa M.H. Ibrahim, Mohammed Nadim Sardoiwala, Megan M. Nelson, Jamie Stout, S. Pablo Sardi, Gabriel Corfas

**Affiliations:** Kresge Hearing Research Institute and Department of Otolaryngology - Head and Neck Surgery, University of Michigan, Ann Arbor, MI 48109, USA; F.M. Kirby Neurobiology Center, Boston Children’s Hospital, Boston, MA 02115; Department of Neurology and Otology and Laryngology, Harvard Medical School, Boston, MA 02115

**Author notes:** These authors contributed equally (Mostafa M.H. Ibrahim, Mohammed Nadim Sardoiwala). Corresponding author: Gabriel Corfas, Ph.D., Kresge Hearing Research Institute, The University of Michigan, Medical Sciences I Building, Rm. 5428, 1150 West Medical Center Drive, Ann Arbor, MI 48109-5616. S.P.S.: Sanofi, Cambridge, MA, USA.

**Keywords:** Neuregulin 1, receptor tyrosine kinase, neural precursor cells, GFAP, nuclear signaling

## Abstract

Precise timing of astrogenesis during brain development is essential for proper neural circuit assembly, yet the molecular mechanisms governing this process remain incompletely understood. Here, we identify a novel nuclear signaling pathway initiated by the presenilin-dependent proteolytic release of the ErbB4 receptor intracellular domain (E4ICD) that regulates astrocyte differentiation via direct interaction with Poly(ADP-ribose) Polymerase 1 (PARP1). Upon ligand stimulation, E4ICD forms a complex with PARP1, inducing PARP1 tyrosine phosphorylation and enzymatic activation. This complex translocates to the nucleus, where it binds to the GFAP promoter, repressing its transcription. Loss of ErbB4 or PARP1 abolishes these effects, leading to increased differentiation of neural precursor cells into astrocytes in vitro, and elevated astrocytic gene expression in the neonatal brain in vivo. Notably, the E4ICD binding sites in the GFAP promoter overlap with those of Notch signaling, suggesting that these signaling pathways have antagonistic transcriptional roles in the control of glial fate. These findings establish a previously unappreciated RTK–PARP1 nuclear signaling axis that integrates extracellular cues to regulate neurodevelopmental lineage decisions, expanding the known functional repertoire of PARP1 beyond DNA damage and stress responses.

## Introduction

Receptor Tyrosine Kinases (RTKs) constitute a family of transmembrane signaling molecules that participate in many key cellular processes, including proliferation, survival, migration, and differentiation [1]. RTKs have long been thought to exert their biological effect through what can now be considered canonical RTK signaling. In this signaling modality, ligand binding induces receptor dimerization, resulting in receptor trans-phosphorylation that creates binding sites for adaptor proteins that then mediate activation of downstream soluble kinases that go on to phosphorylate targets such as transcription factors, which then alter transcription of their specific target genes [2, 3]. Studying ErbB4, a receptor for neuregulin-1 (NRG1) and other trophic factors, we and others discovered a novel mechanism for RTK function involving direct nuclear signaling. In this alternate RTK pathway, ligand-induced activation of the ErbB4-JMa isoform induces cleavage of the receptor, first by the tumor necrosis factor-α-converting enzyme (TACE) in the extracellular juxtamembrane domain [4, 5], then by γ-secretase within the transmembrane domain [6]. This sequence of events results in the release of the activated soluble intracellular domain (E4ICD), which has a nuclear localization sequence that mediates its translocation to the nucleus [6, 7].

There is mounting evidence that soluble E4ICD plays an important role in transcriptional regulation through interactions with a variety of binding partners across multiple tissues. For example, in mammary tissue, E4ICD has been shown to interact with STAT5a to drive expression of β-casein [8], and its interaction with estrogen receptor increases proliferation rates in breast cancer [9]. In lung, E4ICD has been shown to interact with YAP to promote cellular maturation [10]. In the embryonic brain, we showed that E4ICD influences neuronal precursor fate specification by suppressing astrocytic genes via an interaction with the adaptor protein Tab2 and transcriptional coregulatory protein N-CoR [11]. In each of these cases, E4ICD interacts with other nuclear proteins that influence transcription through protein-protein and protein-DNA interactions. However, how E4ICD alters the activity of its partners and whether it induces the activation of other nuclear enzymes remains unclear.

Here we identify the multifunctional enzyme Poly(ADP–ribose) Polymerase 1 (PARP1) as a binding partner for E4ICD and demonstrate that the E4ICD/PARP1 interaction is necessary for the ability of NRG1 to repress the expression of astrocyte genes by neural precursor cells (NPCs) in culture. Furthermore, we show that loss of PARP1 function phenocopies the increased GFAP expression observed in the brains of neonatal ErbB4^-/-^ mice, indicating that NRG1/E4ICD/PARP1 signaling regulates astrogenesis during the late stages of embryonic development. Moreover, we found that active E4ICD phosphorylates PARP1 in tyrosine 907 (Y907), an event associated with increased PARP1 enzymatic activity and reduced binding to certain PARP1 inhibitors [12, 13]. Together, these results provide new insights into how ErbB4 nuclear signaling regulates transcription in neural cells and identify new roles for PARP1 in brain development.

## Results

### Activated E4ICD interacts with PARP1 in yeast and mammalian cells

To gain further insights into the mechanisms by which ErbB4 nuclear signaling regulates transcription, we screened for E4ICD binding proteins in mammalian cells using LexA-E4ICD, a 102 kDa fusion between the human E4ICD and the LexA dimerization domain [11], as bait. LexA mediated dimerization leads to E4ICD activation and auto-phosphorylation, and its expression in murine NPCs leads to the same biological effect as the activation of endogenous ErbB4-JMa [11]. We expressed LexA-E4ICD in HEK293 cells, performed immunoprecipitation with an ErbB4 antibody and subjected samples to mass spectrometric (MS) analysis. This approach identified Poly (ADP-ribose) polymerase 1 (PARP1) as a potential E4ICD interacting protein (20 peptides identified, 36% coverage, see Fig. 1A). PARP1 is a nuclear enzyme that catalyzes the transfer of adenosine diphosphate-ribose (ADP-ribose) units from molecules of nicotinamide adenine dinucleotide (NAD+) to acceptor Glu, Asp, and Lys residues in target proteins, i.e. PARylation, creating highly negatively charged polymers (PAR groups) [14]. PARP1 is involved in numerous key cellular processes, including DNA repair, chromatin remodeling, and transcription [15–18].

**Figure 1.**
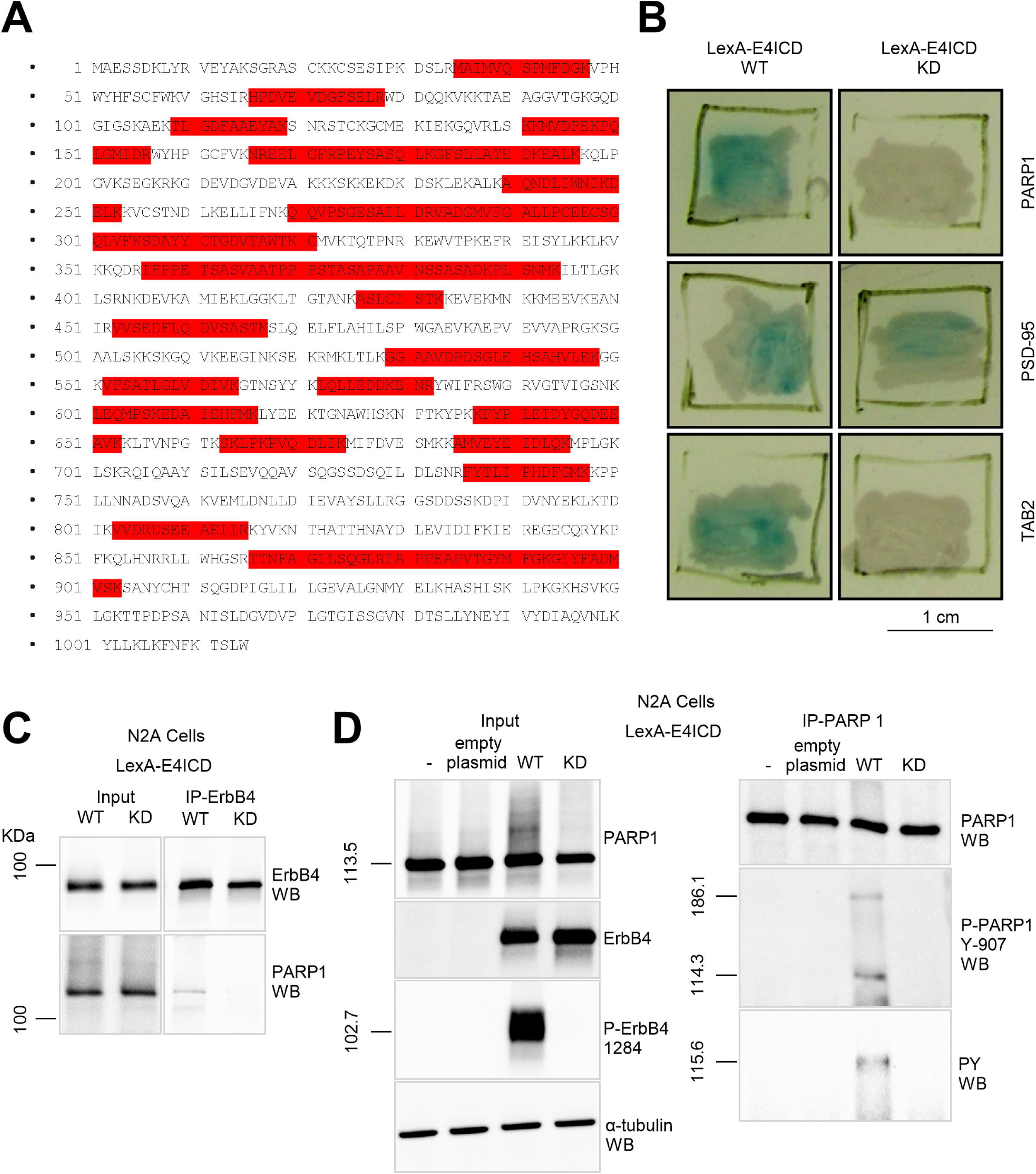
Identification of PARP1 as a LexA-E4ICD interacting partner. **A.** The amino acid sequence of PARP1 with peptides (highlighted in red) identified by mass spectrometry from proteins co-precipitating with LexA-E4ICD. **B.** Yeast-2-Hybrid assay shows that PARP1 (prey) interacts with a bait encoding wild type (WT) LexA-E4ICD (bait) but not a LexA-E4ICD kinase-dead mutant (KD). As controls, consistent with Sardi et al. (2006) [11], PSD95 interacts with E4ICD independently of its kinase activity, while TAB2 requires kinase activity for interaction. Blue colonies indicate positive bait-prey interactions. **C.** Immunoprecipitation with ErbB4 antibodies shows that the active LexA-E4ICD but not the kinase dead (KD) co-precipitates endogenous PARP1 in N2A cells. **D.** Immunoprecipitation with PARP1 antibodies in N2A cell lysates transfected with WT or KD LexA-E4ICD show that PARP1 was only phosphorylated with WT LexA-E4ICD.

To validate the E4ICD/PARP1 interaction, we first used the yeast two-hybrid assay, an approach we had previously utilized to identify and characterize another LexA-E4ICD partner [11]. This approach showed that PARP1, like we previously showed for TAB2 [11], interacts with wild type LexA-E4ICD but not with a kinase-dead version. In contrast, and as we showed earlier [11], E4ICD interaction with PSD95 was unaffected by loss of kinase activity (Fig. 1B). These results support the conclusion that the LexA-E4ICD/PARP1 depends on E4ICD’s kinase activity. Furthermore, co-immunoprecipitation experiments using N2A cells transfected with wild type or kinase dead LexA-E4ICD confirmed that binding of endogenous mammalian PARP1 to E4ICD depends on E4ICD’s tyrosine kinase activity (Fig. 1C). Moreover, wild type LexA-E4ICD expression led to PARP1 tyrosine phosphorylation in residue Y-907 whereas the kinase dead version did not (Fig. 1D). Since this PARP1 phosphorylation event has been linked to PARP1 activation [12], these results suggested that the activated E4ICD complexes with PARP1, leading to its phosphorylation and activation.

### NRG1 induces ErbB4/PARP1 interaction and PARP1 activation in mammalian cells, and this depends on ErbB4-JMa cleavage

Since E4ICD is generated after the cleavable ErbB-JMa isoform is activated by its cognate ligand NRG1, we tested if PARP1 interacts with the full-length ErbB4-JMa in transfected N2A cells, using cells expressing the uncleavable ErbB4-JMb isoform [4, 5] as controls. NRG1 treatment induced ErbB4/PARP1 co-precipitation only in cells expressing ErbB4-JMa (Fig. 2A). Furthermore, PAR Western blot showed that NRG1 induces PARylation only through the cleavable ErbB4-JMa, and that this activity is blocked by TACE or γ-secretase inhibition (Fig. 2B). These results indicate the NRG1 induces PARP1 activity and PARylation through the release of E4ICD after ligand-induced ErbB4-JMa activation.

**Figure 2.**
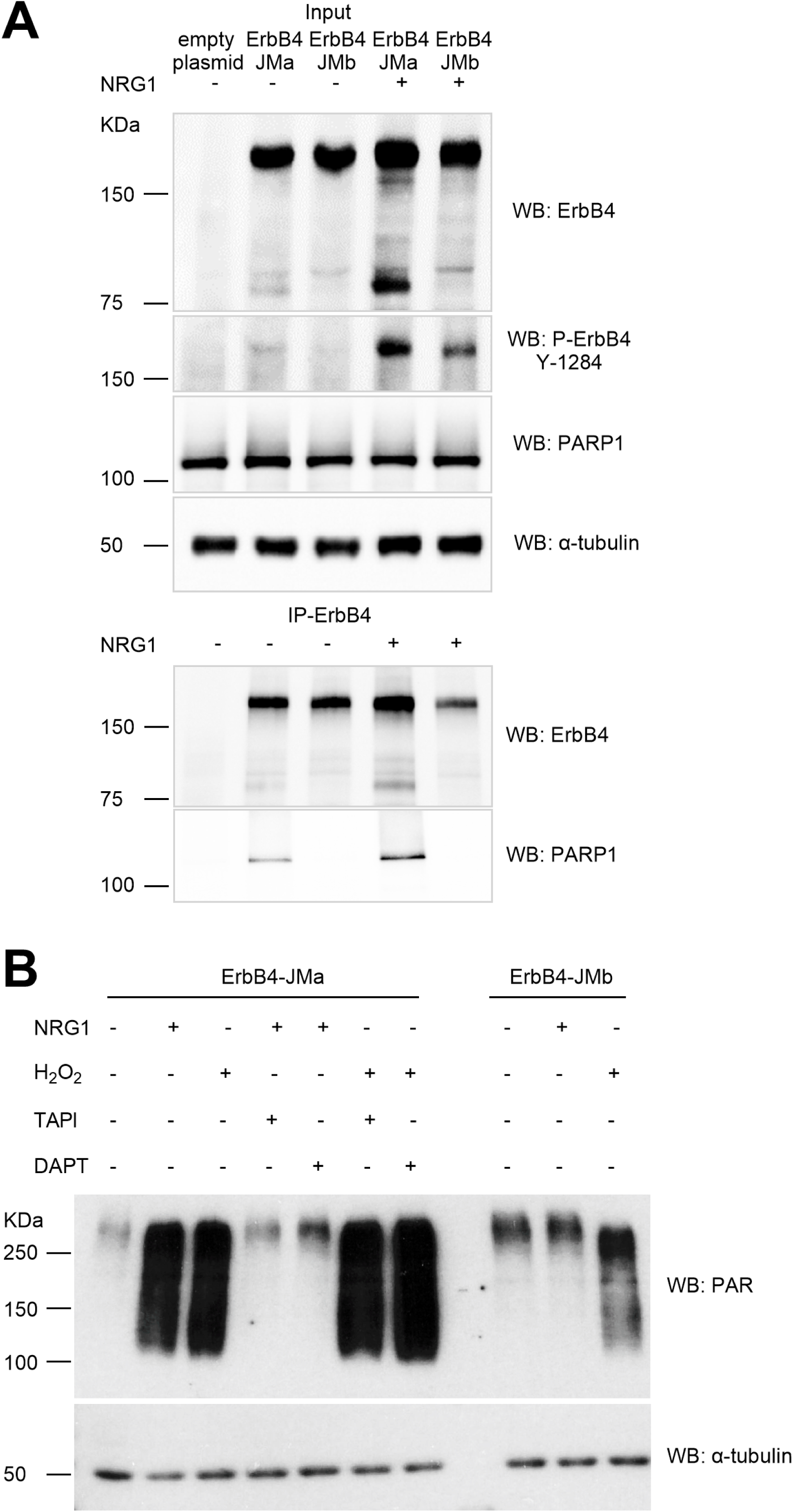
E4ICD binds to PARP1 after NRG1-dependent cleavage of ErbB4-JMa. **A.** NRG1 treatment induces co-precipitation of PARP1 and ErbB4 in N2A cells when cells express the cleavable (ErbB4-JMa) but not the uncleavable (ErbB4-JMb) isoform. **B.** PAR Western blot of lysates of N2A cells treated with 2nM NRG1 or 100μM H_2_O_2_ for 5 min shows that both induce similar shifts in the PARP1 bands and PARylation. Blocking TACE (100μM TAPI) or PS1 (1μM DAPT) inhibits the effects of NRG1 but not of H_2_O_2_ in ErbB4-JMa and not ErbB4-JMb transfected cells where NRG-1 treatment failed to induce PARP1 activation.

We previously demonstrated that ErbB4-JMa nuclear signaling represses the differentiation of rat NPCs into astrocytes [11]. Therefore, to test the biological relevance of the ErbB4-JMa-PARP1 interaction, we used cells and tissues from ErbB4 wild type and KO mice. As shown in Fig. 3A, PARP1 co-precipitated with ErbB4 only in wildtype cells, and NRG1 treatment increased the extent of the co-precipitation. Furthermore, PARP1 co-precipitated with ErbB4 from lysates of E14.5 wild type brains but not from KO tissues (Fig. 3B). These results indicate that ErbB4 has a low level of activation in untreated cells, that exogenous NRG1 increases it, resulting in increased E4ICD/PARP1 interaction, and that E4ICD/PARP1 interact in the intact embryonic brain.

**Figure 3.**
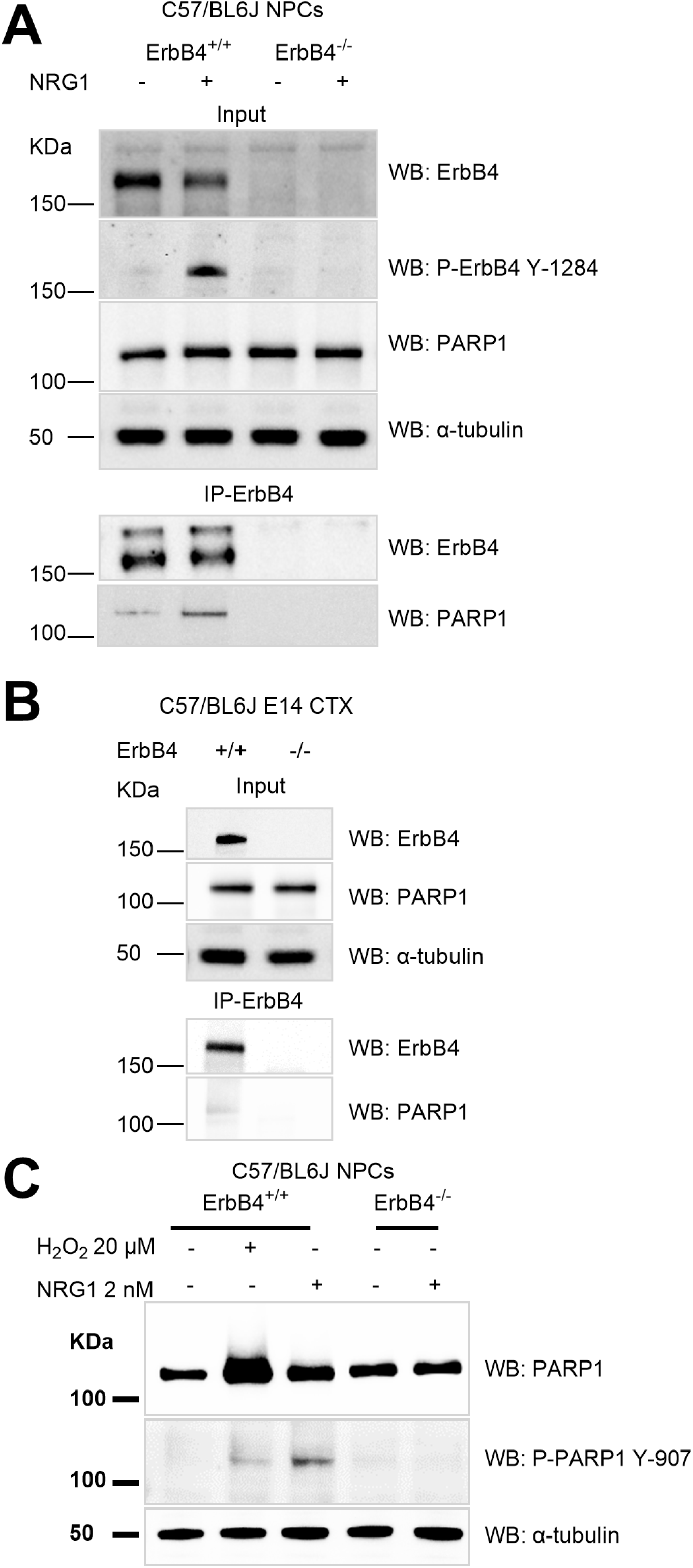
NRG1 induces ErbB4/PARP1 interaction and PARP1 activation in mammalian cells and in vivo. **A.** NRG1 treatment induces co-precipitation of endogenous PARP1 with endogenous ErbB4 in mouse primary neural precursor cells (NPCs). **B.** ErbB4 antibodies co-precipitate PARP1 from extracts of E14.5 wild type mouse brain but not from ErbB4 KO tissues. **C.** Phospho-PARP1 western blot analysis of WT and E4KO NPCs show that 6 hours NRG1 (2nM) and 30 minutes H_2_O_2_ (20 µM) treatment of ErbB4 induces PARP1 phosphorylation in only ErbB4 WT and not ErbB4 KO NPCs.

To determine the E4ICD/PARP1 interaction in neural cells has a functional consequence, we tested whether NRG1 treatment influences PARP1 activity (PARylation) and phosphorylation in NPCs as we observed in cells transfected with ErbB4-JMa, and if this depends on ErbB4 nuclear signaling. Exposure to H_2_O_2_, a well-known PARP1 activator [12, 19, 20], was used as a positive control. In agreement with what we found in N2A cells expressing LexA-E4ICD (Fig. 1 D), NRG1 treatment induced PARP1 Y907 phosphorylation in wild type but not ErbB4^-/-^ NPCs (Fig. 3C). Furthermore, PAR immunostaining showed that NRG1 and H_2_O_2_ treatments induce PARylation in wild type NPCs, but NRG1 fails to do so in ErbB4^-/-^ NPCs, whereas both NRG1 and H_2_O_2_ fail to induce PARylation in PARP1^-/-^ NPCs (Fig. 4). Importantly, EGF, which activates the EGFR (ErbB1) but not ErbB4 [21], does not induce PARylation in NPCs (Fig. 5). Since NPCs express EGFR and respond to EGF [22], these results indicate that the ErbB4-mediated PARP1 activation in NPCs is specific rather than a consequence of all RTK signaling. Interestingly, the pattern of PARylation in NPCs following NRG1 and H_2_O_2_ treatments were significantly different. Whereas H_2_O_2_ induces PARylation rapidly and transiently in both nuclei and cytoplasm, the NRG1-induced PARylation starts in the cytoplasm within 5 minutes of treatment and gradually moves to the nuclei (Fig 6). These results indicate that ligand-induced ErbB4-JMa signaling activates PARP1, which leads to a wave of PARylation that moves from the cell cytoplasm to the nucleus.

**Figure 4.**
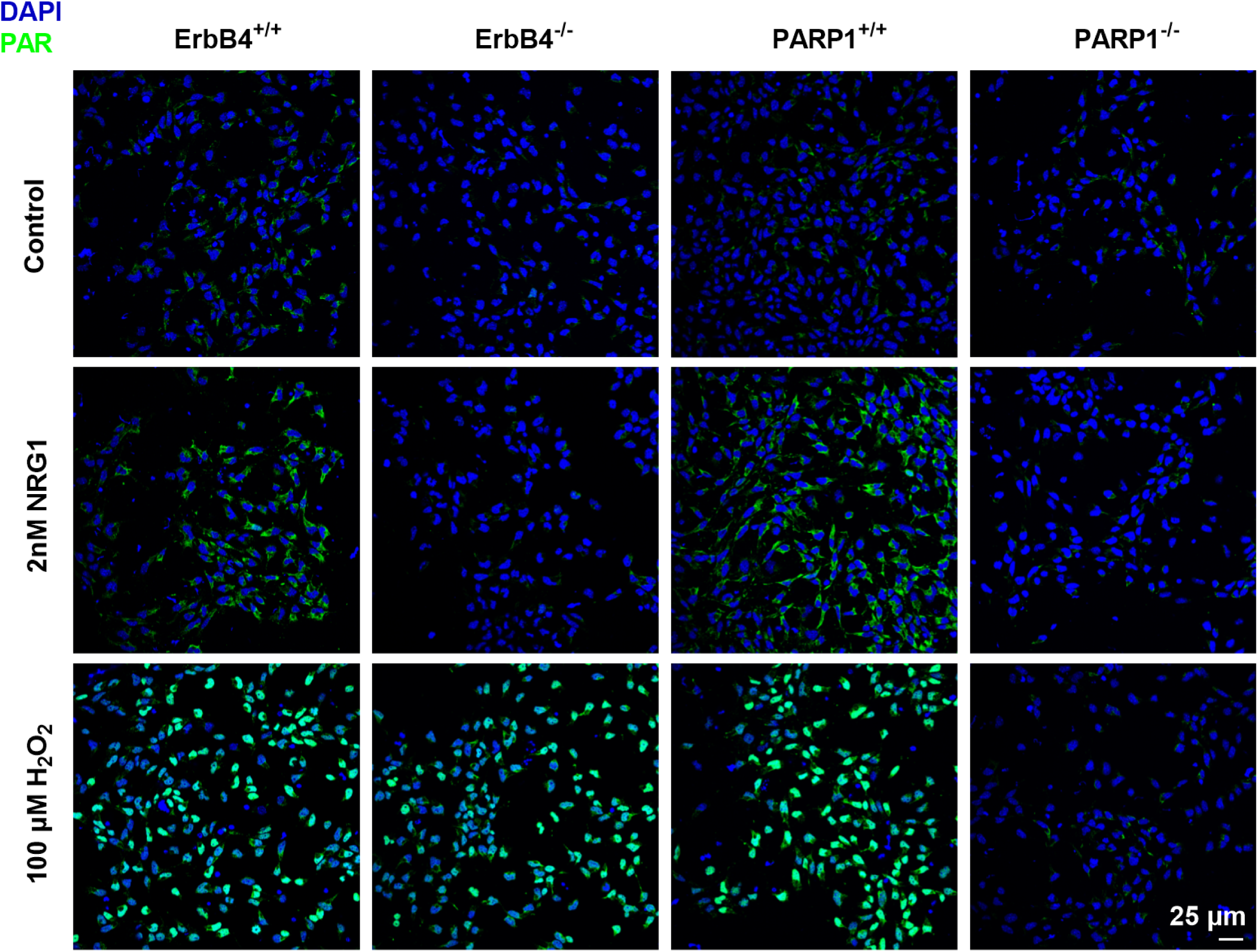
ErbB4 and PARP1 are necessary for NRG1-induced PARylation in NPCs. Immunostaining for PAR in NPCs shows an upregulation of PARylation upon 5 min treatment with NRG1 (2nM) or H_2_O_2_ (100μM) in WT NPCs. NRG1-induced PARylation is absent in ErbB4 KO and PARP1 KO NPCs, while neither NRG1 nor H_2_O_2_ induce PARylation in PARP1 KO NPCs.

**Figure 5.**
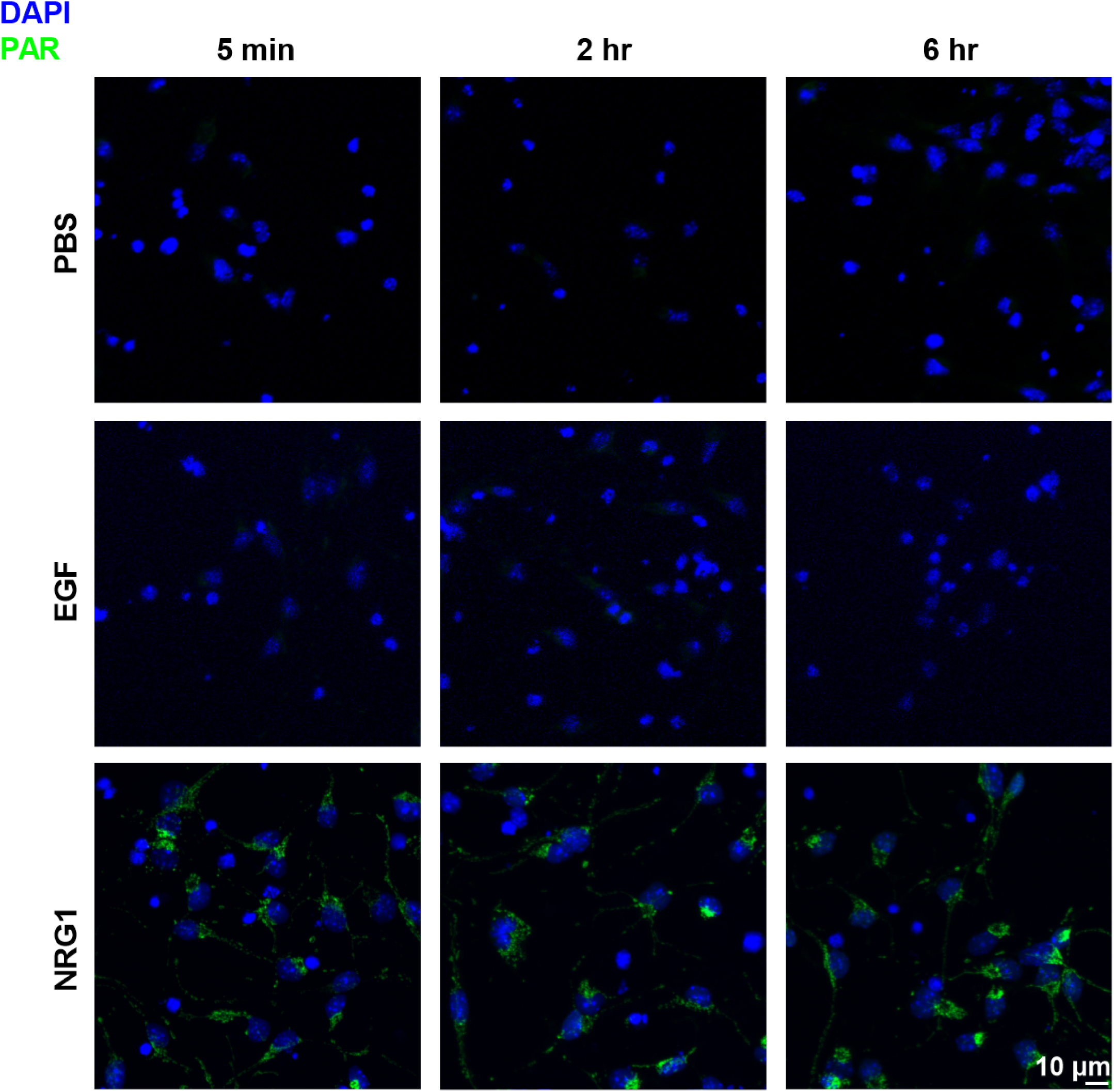
PARylation in NPCs is induced by NRG1 but not EGF. Immunostaining for PAR in WT NPCs shows an upregulation of PARylation upon 5 min treatment with NRG1 (2nM) but not EGF (20ng/mL).

**Figure 6.**
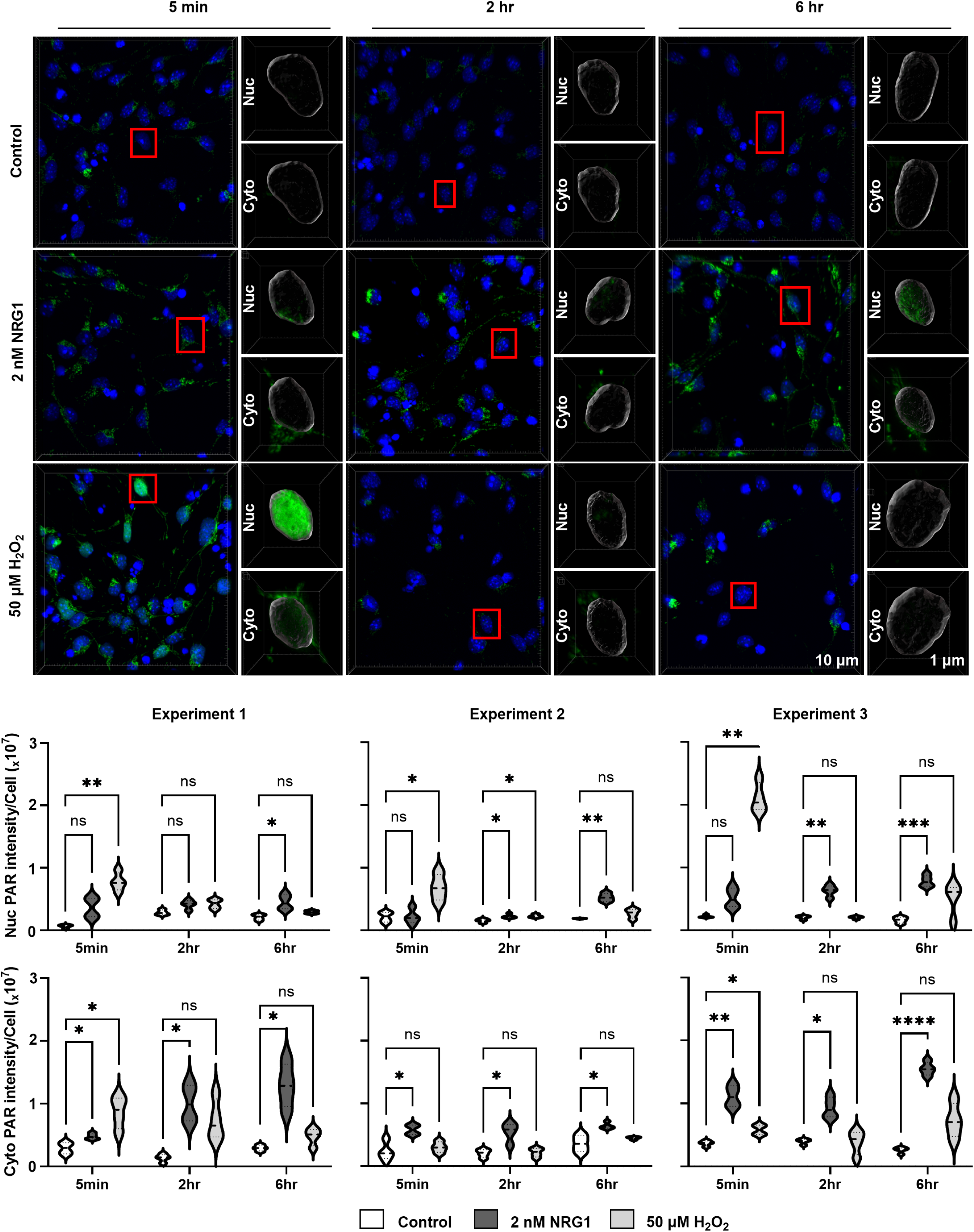
NRG1 and H_2_O_2_ show different patterns for PARylation in NPCs. <u>TOP:</u> PAR immunostaining of NPCs treated with either PBS (control), NRG1 (2nM) or H_2_O_2_ (50μM), for 5 minutes, washed and either fixed immediately (5 min), or incubated for an addition 2 or 6 hours. The insets depict representative staining in nuclei (Nuc) and perinuclear cytoplasm (Cyto). <u>BOTTOM:</u> quantitative analysis of 3 independent experiments shows that NRG1 (2nM) treatment leads to rapid cytoplasmic PARylation which then moves to the nucleus 2-6 hours later. In contrast, H_2_O_2_ (50μM) induces nuclear and cytoplasmic PARylation rapidly and transiently.

### NRG1-dependent regulation of astrocyte gene expression in NPCs requires ErbB4-JMa and PARP1

We previously reported that NRG1 represses the differentiation of rat and mouse NPCs into GFAP-expressing cells through ErbB4 nuclear signaling, and that this depends on ErbB4-JMa [11, 23]. To test if this effect of NRG1-ErbB4-JMa signaling requires PARP1, we used NPCs derived from ErbB4^-/-^, ErbB4-JMa^-/-^, or PARP1^-/-^ E14.5 mouse embryos and their respective wild types. As we showed before, NRG1 treatment (2 nM) led to a significant reduction in GFAP and S100β mRNA levels in wild type NPCs (ErbB4^+/+^ and PARP1^+/+^) after bFGF removal but fail to do so in all the knockout models (Fig. 7). Importantly, the levels of expression of markers of NPCs (Nestin [24]), neurons (TUJ1/β3 tubulin [25–27]), and oligodendrocytes and their precursors (Olig2 [28]) were not influenced by NRG1 treatment in any case, indicating that NRG1/ErbB4-JMa/PARP1 signaling might be critical for astrogenesis but not for the generation of other key brain cell types.

**Figure 7.**
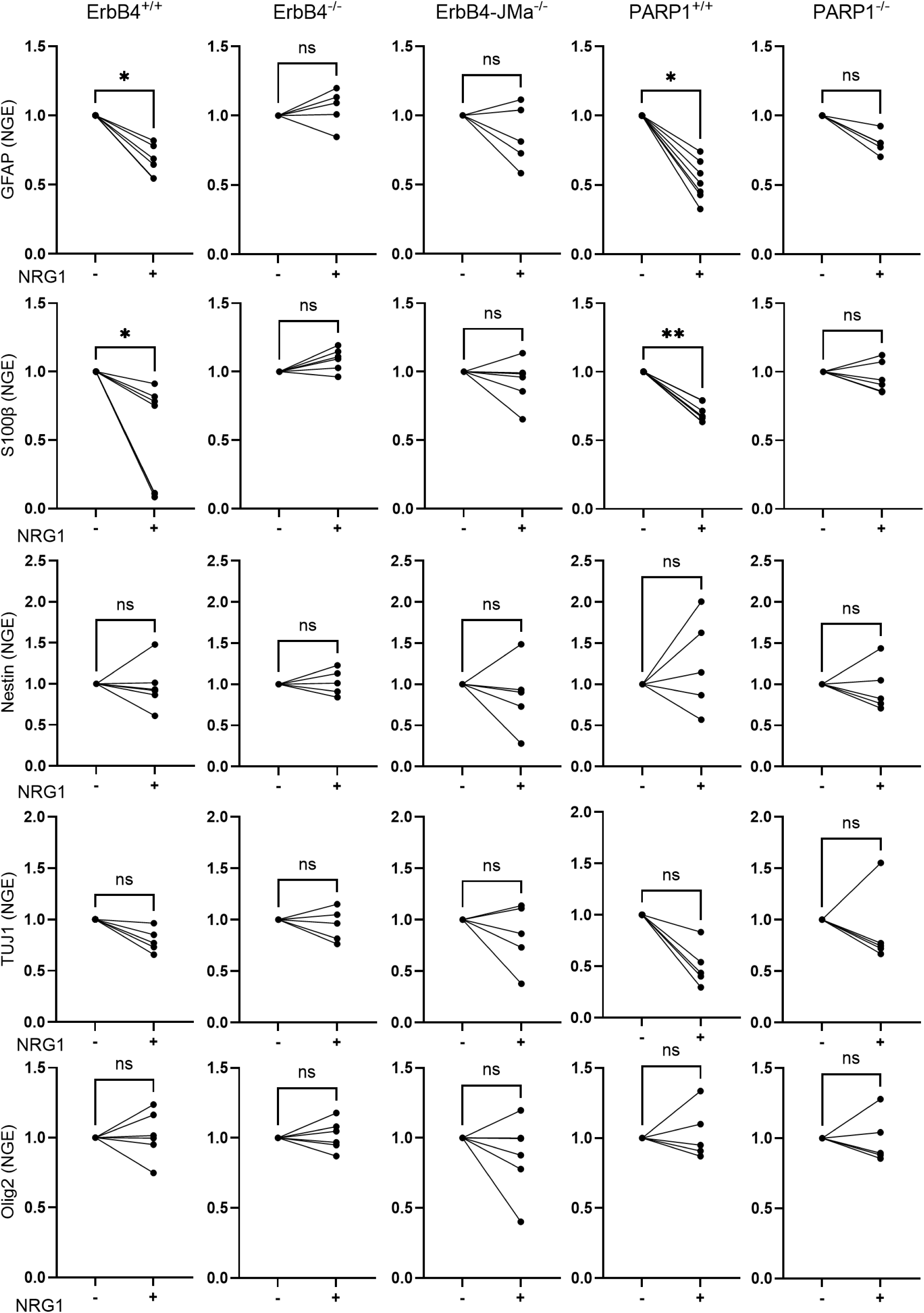
E4ICD/PARP1 Interaction Represses Astrocytic Gene Expression. RT-qPCR shows that GFAP and S100β mRNA expression are significantly reduced in WT NPCs treated with NRG-1 compared to untreated WT NPCs. No significant differences in GFAP or S100β transcripts were observed in all KOs NPCs (ErbB4^-/-^, JMa^-/-^ and PARP1^-/-^), with or without NRG-1 treatment. No significant differences in Nestin, TUJ1 or Olig2 transcripts were observed in WT or KO NPCs with or without NRG-1 treatment; n = 5 per group. Paired Student’s t-test was used to evaluate differences. mRNA Expression levels were quantified relative to RPL19.

### Loss of ErbB4 and PARP1 function leads to similar increase in GFAP expression in the neonatal brain

To determine if ErbB4 and PARP1 regulate astrocyte gene expression *in vivo*, we measured the mRNA expression levels of three astrocytic markers (GFAP, S100β and ALDH1L1) in the brain of ErbB4^-/-^ and PARP1^-/-^ newborn mice and their respective wild types. Loss of ErbB4 or PARP1 led to increased mRNA levels for the three genes (Fig 8A). Consistent with those findings, there was a significant increase in GFAP protein levels in the brains of both ErbB4 and PARP1 knock out mice. Interestingly, the level of FABP7 expression, a radial glial cells marker [29], was not altered in the mutants, indicating that the main effect of loss of ErbB4 and PARP1 is on astrogenesis (Figs. 8B and C). Furthermore, histology showed a clear elevation of GFAP immunostaining in the hippocampus of newborn KOs compared to the wild types (Fig. 9). Together, these results indicate that both ErbB4 and PARP1 are necessary for the repression of astrogenesis in the late stages of embryonic brain development, and the loss of either one of them disrupts the NRG1-ErbB4 signaling-dependent regulation of astrogenesis.

**Figure 8.**
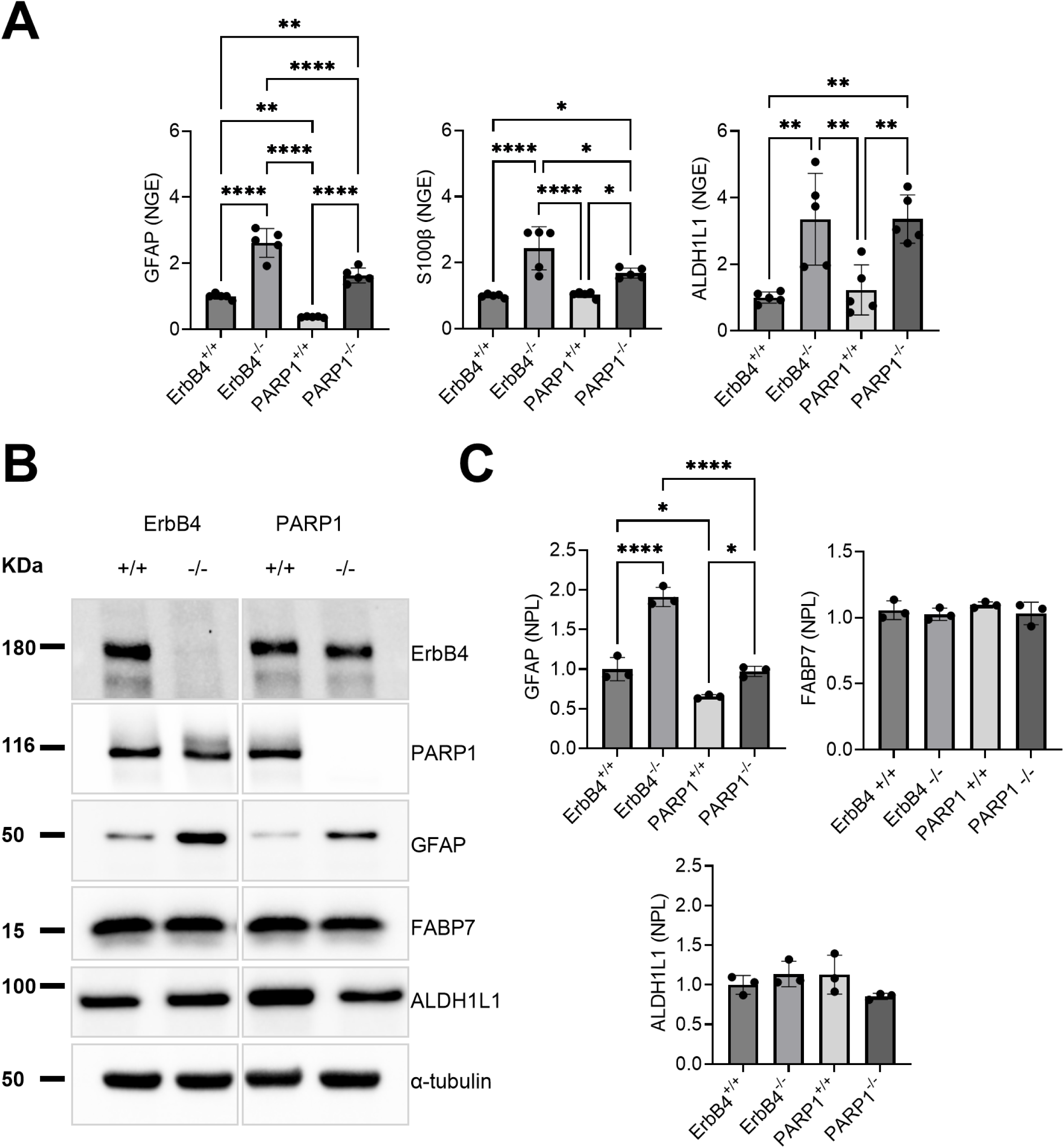
ErbB4 or PARP1 knock out Induces Increase in Astrocytic Markers in Neonatal Pups. **A.** RT-qPCR shows that GFAP, S100β and ALDH1L1 mRNA expression levels are significantly higher in ErbB4 and PARP1 KO P0 mouse brain tissues compared to their WT counterparts; n = 5 per group. One way ANOVA was used for analysis. **B.** GFAP western blot analysis of P0 cortical tissue of WT, PARP1 KO and ErbB4 KO mice shows mutants have much higher levels of GFAP protein expression than WTs. There is no significant difference in FABP7 or ALDH1L1 protein expression levels when comparing WT to KO tissues. **C.** Quantitative Western blot analysis of GFAP shows that GFAP protein levels increase in KO P0 mouse cortical tissue compared to WT; n = 3 per group. One way ANOVA was used for analysis.

**Figure 9.**
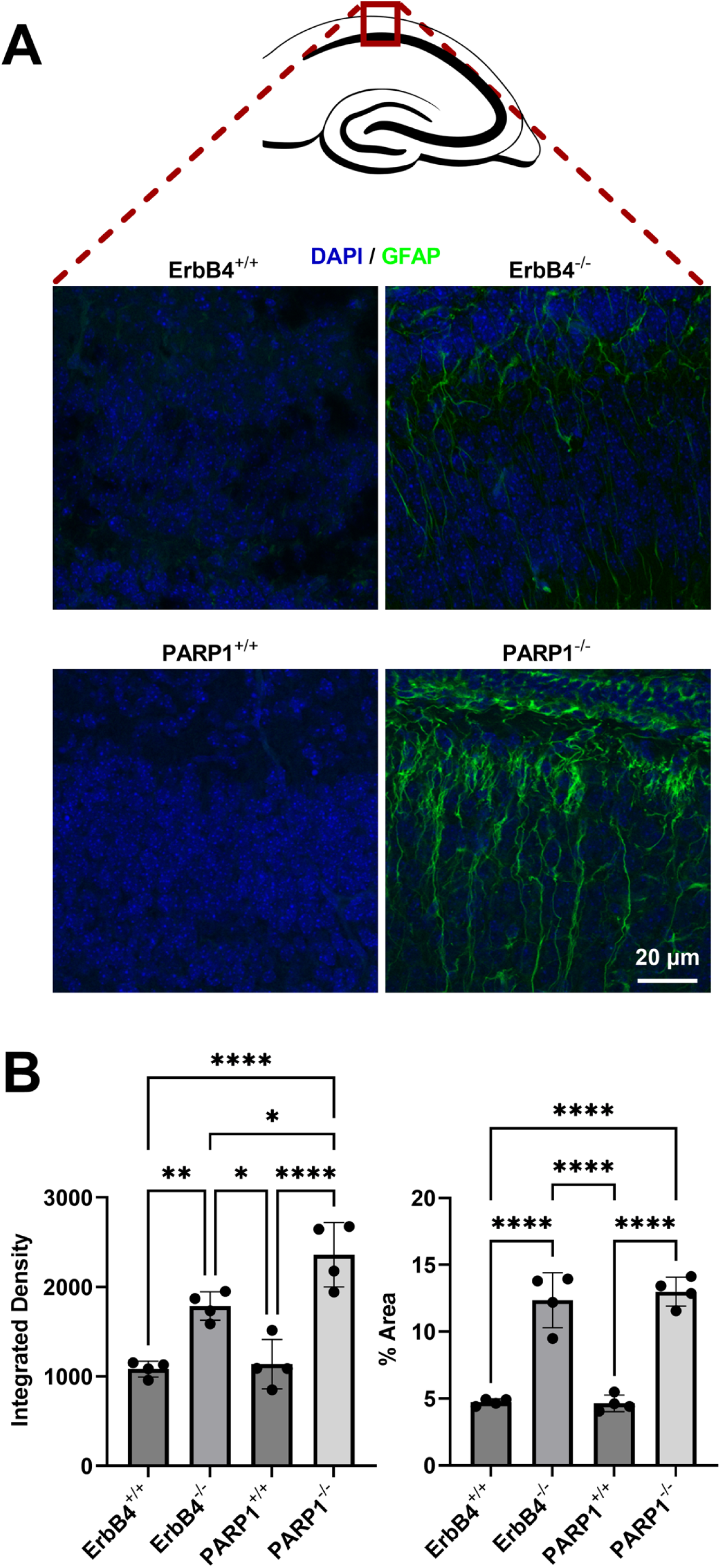
GFAP immunostaining of P0 brain sections of the hippocampus shows that knocking out of either ErbB4 or PARP1 leads to a significant increase in GFAP levels compared to the wild types. There is an increase in both the integrated density and the area percentage indicating an increase in GFAP expression in each cell and a possible increase in the number of astrocytes expressing GFAP; n = 4 per group. One way ANOVA was used for analysis.

### E4ICD binds to GFAP promoter region

We previously showed that E4ICD binds to the GFAP promoter in rat NPCs, suggesting that this could be important for its ability to repress GFAP expression in the developing brain [11].

Interestingly, the genomic sequence we identified as binding E4ICD has been shown to act as a binding site for Recombination signal Binding Protein for immunoglobulin kappa J region (RBPJ), a protein involved in the Notch signaling pathway [30] Binding of the complex formed by RBPJ and the Notch intracellular domain (NICD) to the GFAP promoter activates its expression [30]. To determine if E4ICD binds to some or all RBPJ sites in the GFAP promoter, and if this depends on PARP1, we performed ChIP-qPCR using primers specific for proven [30] and predicted RBPJ[31] binding sites in the mouse GFAP promoter using wild type and ErbB4 KO NPCs that were treated with either NRG1 or vehicle for 6 hours. NRG1 treatment induced E4ICD binding to 5 of the 6 regions tested, an effect that was lost in ErbB4^-/-^ NPCs but not in PARP1^-/-^ cells (Fig.10). Together, these results indicate that NRG1-mediated E4ICD generation leads to the binding of E4ICD to several RBPJ sites in the GFAP promoter, and that this might be critical for its ability to repress GFAP expression in the later stages of embryonic development. Furthermore, since in PARP1^-/-^ cells, E4ICD NRG1-induced E4ICD binding to the GFAP promoter is still present, but the NRG1-induced GFAP repression is lost, the results indicate that PARP1 is necessary for the repressive activity of the E4ICD complex, but not for its ability to bind chromatin.

**Figure 10.**
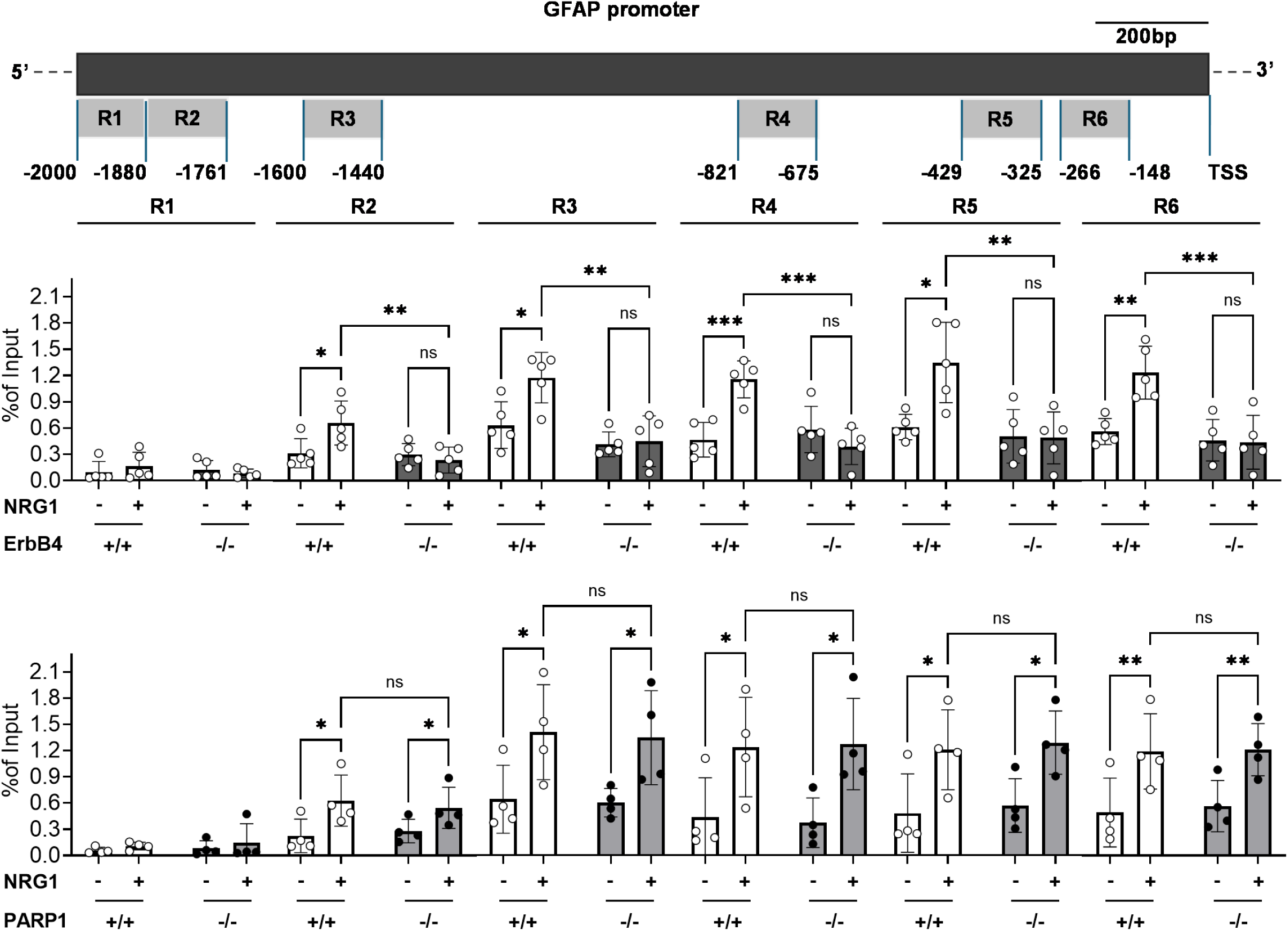
The E4ICD binding assessment to the GFAP promoter regions indicates NRG1 treatment is required for E4ICD binding and PARP1 is not necessary for the E4ICD binding. (Two-way ANOVA, *p<0.05, **p<0.01, ***p<0.001)

## Discussion

Our study identifies PARP1 as a critical mediator of ErbB4 nuclear signaling, acting downstream of ligand-dependent ErbB4-JMa cleavage to regulate astrocyte gene expression in embryonic NPCs. Mass spectrometry of HEK293 cell lysates expressing activated LexA-E4ICD revealed PARP1 as a high-confidence interacting partner. This interaction, confirmed by yeast two-hybrid and co-immunoprecipitation assays, depends on E4ICD’s intrinsic kinase activity mirroring the activation mechanism that drives E4ICD’s repressive function in NPCs. This ligand-activated, proteolysis-dependent, ErbB4-PARP1 signaling axis underscores a novel developmental function for PARP1 as a nuclear coregulator directly engaged by an RTK intracellular domain.

PARP1 is a multifunctional chromatin-associated enzyme with widely documented roles in transcriptional regulation, including modulation of chromatin architecture, enhancer binding, and coregulator functions that can activate or repress gene expression depending on context [17, 18, 32–34]. PARP1 activities include orchestrating nucleosome remodeling, regulating histone composition, and facilitating dynamic cofactor exchange at target promoters [18, 33, 35, 36]. Our data extends this well-established framework by characterizing a specialized developmentally regulated pathway in which PARP1 is recruited and tyrosine phosphorylated by E4ICD for transcriptional repression of astrocyte genes during neurogenesis. Unlike the rapid nuclear PARylation typically triggered by genotoxic stress, ErbB4 signaling induces a phased and spatially regulated PARP1 activation beginning in the cytoplasm and culminating in chromatin repression. This highlights how PARP1’s diverse transcriptional regulatory mechanisms can be tailored to developmental signaling pathways distinct from hormone responses or cellular stress.

A key mechanistic insight from our study is that E4ICD not only binds PARP1 but also induces its phosphorylation on tyrosine 907 (Y907), a post-translational modification known to enhance enzymatic activity and reduce affinity for PARP inhibitors [12, 13, 37, 38]. Similar phosphorylation events have been described downstream of other receptor tyrosine kinases, including EGFR and MET (c-Met), where PARP1 Y907 phosphorylation supports DNA repair and survival signaling, particularly under oxidative stress and in cancer contexts [12, 39, 40]. Notably, in our experiments, NRG1-induced PARP1 phosphorylation depended on ErbB4-JMa expression and was absent in ErbB4-null NPCs, indicating that ErbB4, like EGFR and MET, serves as an upstream RTK regulator of PARP1 activity. However, unlike these other RTKs, ErbB4-induced phosphorylation of PARP1 occurs in a physiological, developmental context and is functionally coupled to transcriptional repression of astrocyte-specific genes rather than DNA repair or pro-survival signaling. Moreover, EGFR activation by EGF failed to induce PARP1 activation in NPCs, highlighting the signaling specificity and distinct biological outcome of ErbB4-PARP1 nuclear signaling.

Our ChIP-qPCR analyses show E4ICD binds multiple RBPJ motifs within the murine GFAP promoter in an NRG1- and ErbB4-dependent manner, overlapping with binding sites for the Notch intracellular domain (NICD), a key activator of astrocytic gene expression. Importantly, loss of PARP1 abolishes E4ICD-mediated repression without affecting E4ICD chromatin occupancy, indicating that PARP1 functions as a critical corepressor within this complex. This highlights a potential antagonistic transcriptional regulation at shared genomic loci by ErbB4 and Notch signaling pathways, adding a new dimension to how PARP1-regulated chromatin remodeling interfaces with lineage decision networks.

Loss of ErbB4 or PARP1 results in failure of NRG1 to repress astrocytic markers in NPC culture and leads to elevated astrocyte gene expression and protein levels, notably GFAP, in neonatal hippocampus in vivo. These effects are specific, sparing neuronal and oligodendrocyte lineage markers, consistent with a key checkpoint role for ErbB4-PARP1 signaling in timing astrocyte differentiation. Given the important functions of astrocytes in synaptic maturation and plasticity, particularly in the hippocampus, aberrant signaling through this axis may have implications for neural circuit development and neuropsychiatric disorders. Moreover, due to the widespread clinical use of PARP inhibitors, our findings raise caution about their potential unintended effects on developmental astrogenesis and brain function.

Together, our study defines a ligand-triggered nuclear RTK signaling mechanism wherein ErbB4 proteolytically releases E4ICD, which complexes with and phosphorylates PARP1, enabling its nuclear translocation and transcriptional repression of astrocyte genes at RBPJ sites. This repressive function competes with Notch–NICD-mediated activation at the same loci, revealing a bifunctional regulatory module critical for neurodevelopmental fate decisions. Our work expands the biological scope of PARP1 from classical stress and DNA repair contexts to include its role as a developmentally regulated corepressor within a proteolytically activated receptor pathway, underscoring PARP1’s functional versatility in transcriptional control.

## Material and methods

### Animals

An ErbB4 knock-out mouse line that is rescued from lethality due to cardiac defects by expressing human ErbB4 cDNA under the cardiac-specific myosin heavy chain (MHC) promoter (*ErbB4−/−*) [41] was kindly provided by Dr. Martin Gassmann (Department of Biomedicine, Institute of Physiology, University of Basel, Switzerland). This line is in a C57BL/6 background. The PARP1 KO mouse line 129S-*Parp1tm1Zqw*/J (Wang et al., 1995) was obtained from the Jackson Laboratory. This line is in a 129S1/SvImJ background. All animals were maintained under a 12/12 h light/dark cycle and received food ad-libitum. Animal procedures were reviewed and approved by the Animal Care and Use Committee and the University of Michigan Institutional Animal Care and Use Committee.

### Animal genotyping

Ear snips less than 3 mm were collected prior to weaning at P21. The DNA was isolated using the Hot-Shot genomic DNA extraction method using alkaline lysis buffer for 30 minutes at 95°C followed by halting the process with neutralization buffer (40 mM Tris-HCl). Isolated DNA was used for PCR assay using GoTaq® Green Master Mix (Promega Cat. No. M712C) which includes Taq DNA polymerase, dNTPs, MgCl_2_ and reaction buffers at optimal concentrations. The primers used are: (ErbB4 Forward primer: CAGTGTGCGCAGGAACAGAGAAC, ErbB4 WT Reverse primer: AGACCGCAGGAAGGAGAGGTC, ErbB4 mutant Reverse primer: CATCTGCACGAGACTAGTGAGAC, MHC Forward primer: AGCTGTGGTCCACATTCTTCAGGA, MHC Reverse primer: ACTTGCGCAAGGCTCGGTACTGCT, PARP1 Forward primer: CATGTTCGATGGGAAAGTCCC, PARP1 WT reverse primer: CCAGCGCAGCTCAGAGAAGCCA and PARP1 mutant reverse primer: AGGTGAGATGACAGGAGATC). The PCR conditions were 94°C for 2 minutes followed by 35 cycles of 30 seconds at 94°C, 30 seconds at 65°C and 30 seconds at 72°C, then lastly 2 minutes at 72°C. The PCR products were run on a 2% agarose gel in 1X TAE buffer.

### Cell Culture

For mouse NPCs, telencephalons were isolated from E14.5 embryos, placed on ice cold PBS, dissociated and expanded in T75 tissue culture flasks containing DMEM supplemented with 2% B27 (Invitrogen), 20 ng/mL epidermal growth factor (EGF), and basic fibroblast growth factor (bFGF; 20 ng/ml), in a humidified 5% CO2/95% air incubator at 37°C. Within 3 days, the cells grew as free-floating neurospheres, which were then dissociated with accutase and plated as monolayer on poly-L-lysine and fibronectin coated plates in DMEM supplemented with 2% B27 and 20 ng/ml bFGF. N2A cell lines were cultured in DMEM supplemented with 10% fetal bovine serum.

### Plasmids and transfections

Full-length cDNAs encoding human ErbB4-JMa and ErbB4-JMb cloned into pCDNA3 were used to transiently express ErbB4 isoforms in N2A cells. Wild type and kinase dead Lexa-E4ICD cloned into pCDNA3 (Sardi et al., 2006) were used to transiently express E4ICD in N2A cells. All transient transfections were made using Lipofectamine 3000 according to the manufacturer’s instructions (Invitrogen).

### Proteomics

Samples were subjected to SDS PAGE, the gel stained with colloidal blue and target gel bands were cut into 1mm slices, destained with 100mM ammonium bicarbonate/acetonitrile (1:1, vol/vol), followed by reduction and alkylation with 20mM Dithiothreitol and 10mM iodoacetoamide, respectively. After an overnight trypsin digestion at 37°C, the tryptic peptides were extracted with 1:2 (vol/vol) 5% formic acid/acetonitrile and desalted by C18 ziptip. Peptides were then injected in an in-house packed (75 µm × 15 cm, 3 µM particle size, pore size 100A, Michrom Bioresources, CA, USA) C18 RP analytical column and eluted with a linear gradient, starting from 5% B to 35% B in 30 min (A, water with 0.1% formic acid; B, acetonitrile with 0.1% formic acid), followed by fast increase to 95% B in 1 min, and retaining 5 min, then to 5% B in 2 min. The column flow rate was set at 400 nL/min, and the electrospray voltage was set to 2.1 kV. A Q Exactive Orbitrap mass spectrometer (Thermo) was operated in the data-dependent mode to switch automatically between MS and MS/MS acquisition. Survey full-scan MS spectra (m/z 350–2000) were acquired with a mass resolution of 70,000, AGC target value of 3e6 and max IT of 20ms, followed by top 10 sequential MS/MS scans. Dynamic exclusion was set to 20-second exclusion duration. For MS/MS, precursor ions were activated using 27% normalized collision energy with the isolation window at 1.6 m/z. All MS/MS spectra were identified by using MASCOT (v.2.3.02). A decoy database containing the reverse sequences was appended to the database to reduce false-positive identification results. The parameters used for database searching were set up as follows: trypsin as the protease with a maximum of two missed cleavages allowed. Carbamidomethylation of cysteine was specified as a fixed modification and oxidation of methionine, deamination of asparagine and glutamine, phosphorylation of serine, threonine and tyrosine were included as variable modifications. The minimum peptide length was specified to be 5 amino acids. The mass error was set to 10 ppm for precursor ions and 20 mmu for fragment ions.

### Yeast Two-Hybrid Assay

LexA-E4ICD bait was generated by cloning the intracellular domain of ErbB4 (E4ICD, amino acid residues 676–1308) into pEG202. LexA carries a dimerization domain, thus allowing the bait tyrosine autophosphorylation when expressed in yeast [11]. Site-directed mutagenesis was used to create the mutant (K751M of full length) kinase-dead LexA-E4ICDKD. cDNA encoding for full length PARP1 (Genebank ID: NM_007415) was subcloned into EcoRI/XhoI in pJG4-5 to generate the prey fusion protein with the B42 activation domain. A cDNA encoding for PDZ domains 1 and 2 of PSD-95 was subcloned into pJG4-5 and used as a positive control. Site-directed mutagenesis was used to create the mutant (K751M 1 of full length) kinase-dead LexA-E4ICDKD. All constructs were verified by DNA sequencing. Two-hybrid assays were performed using the yeast strain EGY48 harboring LacZ and LEU2 reporters, as described in [11]. Three independent experiments were performed.

### Co-immunoprecipitation (Co-IP) and Western Blot

For Co-IPs to explore the effect of NRG1 on ErbB4/PARP1 interaction in culture, cells were pretreated with 2nM NRG1 or PBS for 40 min, washed twice with ice cold PBS, and then lysed in ice cold cell lysis buffer (Cell Signaling Technology 9803S) with Halt™ Protease and Phosphatase Inhibitor Cocktail (ThermoFisher Scientific 78442) and PARG inhibitor (MedChemExpress PDD 00017273), a specific inhibitor of poly (ADP-ribose) glycohydrolase (PARG) to sustain the PAR levels throughout the experiment. For *in vivo* co-IP, telencephalons were isolated and lysed quickly into the ice-cold lysis buffer. Lysates were subjected to sonication followed by centrifugation at 16000 RMP at 4C for 15 min. Supernatant protein concentration was measured and normalized using BCA assay (Pierce) and then incubated with Protein G Magnetic Beads (New England Biolabs S1430S) and ErbB4 antibody (Cell Signaling Tech.) overnight. Immune complexes were washed five times overnight with lysis buffer before resuspending in SDS sample buffer and heating for 5 min at 95 °C. Samples were subjected to SDS-PAGE. Western blotting was performed by standard protocols and developed using ECL reagents. At least three independent replicates were performed for each experiment.

The cerebral cortex tissue was dissected from brains of P0 mice pups and homogenized using a dounce homogenizer size A in RIPA buffer (Sigma-Aldrich R0278) with protease and phosphatase inhibitors (Thermo-Scientific 78446) and PARG inhibitor (MedChemExpress PDD 00017273) and left on ice for 15 minutes followed by sonication using a probe sonicator (3 pulses, 2 seconds each on ice). Then the lysates were vortexed for 30 seconds followed by centrifugation at 15,000 RPM for 5 minutes and the protein concentration in the supernatant was determined and standardized using Pierce BCA Protein Assay Kits (Thermo-Scientific 23227). Then the standardized samples were diluted in 4X Laemmli buffer (Bio-Rad 1610747) with 10% β-mercaptoethanol. Samples were run on 4–15% Mini-PROTEAN® TGX™ Precast Gel (Bio-Rad 4561083) at 25 mA per gel for 1 hour. The protein was then transferred from the gel to the PVDF membrane with 0.45 µm pore size for 30 minutes at 10 volts using Trans-Blot® SD Semi-Dry Transfer Cell (Bio-Rad). The PVDF membranes were then incubated for blocking in 5% bovine serum albumin (Sigma-Aldrich) in TBST buffer (0.2% Tween20 in TBS) for 90 minutes at room temperature. The blocked membranes were then incubated with primary antibody 1:1000 (PARP1 Cell Signaling 9532, Alpha Tubulin Santa Cruz sc-23948) in blocking solution at 4°C overnight. The following day, the blots were washed in TBST buffer and incubated with the HRP-conjugated secondary antibody in blocking buffer (1:3000 goat anti-rabbit Cell Signaling or 1:3000 horse anti-mouse Cell Signaling) for 1 hour at room temperature then washed again in TBST buffer. The blots were then exposed using SuperSignal™ West Femto Maximum Sensitivity Substrate (Thermo-Scientific 34095), images were developed on a Bio-Rad Chemidoc and then images were analyzed using Bio-Rad Image Lab software.

### PAR Immunostaining on NPCs in culture

For PAR immunostaining, NPCs were treated with H_2_O_2_ (5 minutes, 100 µM, for figure 4, 50 µM for figure 6), NRG1 (5, minutes, 2nM), EGF (5 minutes, 20ng/mL) or PBS. For the 2- and 6-hour time points, cells were washed with fresh medium and incubated for the remaining of the experiment. At the end point, cells were rinsed with PBS, fixed in 4% paraformaldehyde, blocked in PBS containing 0.2% Triton X-100 and 10% normal donkey serum for 2 hours and incubated overnight with PAR antibody diluted in blocking buffer at 1:500. Labeling was visualized using Alexa Fluor secondary antibodies in conjunction with nuclear staining with DAPI. Imaging was performed using a Leica SP8 confocal.

### P0 Cortex immunofluorescence

Whole brains were dissected from P0 pups in the afternoon following birth and fixed in 4% PFA at 4 °C overnight and then cryopreserved in 30% sucrose in PBS for 2 days at 4 °C. Brains were embedded in OCT Compound (Fisher) and snap frozen in isopentane on dry ice. Frozen brains were cryosectioned at 50 μm into antifreeze solution and stored at −20 °C. Sections were washed/permeabilized in TBS 0.2% Triton (TBS-T) for 20 min and blocked in 5% normal donkey serum in TBS-T for 1 h. Sections were then incubated overnight at 4 °C in primary antibody (1:1000. GFAP, (E4L7M) XP® Rabbit mAb #80788, Cell Signaling) then washed in TBS-T at room temperature. Sections were incubated in secondary antibody (1:1000, Alexa fluor 488 nm donkey anti-rabbit, Invitrogen) for 1 h at room temperature then washed in TBS-T and coverslipped with Fluoro-Gel II with DAPI (Electron Microscopy Services) on Superfrost plus slides (Fisher).

### RNA extraction and quantitative RT-PCR

NPCs were grown into neurospheres and then plated at 1×10^6^ cells per well in 6-well plates with bFGF. The second day, bFGF was replenished to prevent differentiation. The third day, all the media was changed without bFGF and wells were treated with 2 nM NRG1 (R&D Systems) or vehicle. After 24 h, total RNA was extracted according to RNeasy Mini Kit (Qiagen) for qRT-PCR. Brains were harvested from newborn pups (P0) and the forebrains were dissected followed by RNA extraction with RNeasy Protect Mini Kit (250) (Qiagen Cat. No. 74126). Following the manufacturer’s instructions. DNase digestion was done on column using RNase-Free DNase kit (Qiagen Cat. No. 79256). RNA quantification and quality assessment was performed on a BioTek plate reader using a BioTek Take3 Trio Microvolume Plate. Equal amounts of RNA per sample (500 ng) were reverse transcribed into cDNA using iScript cDNA Synthesis Kit (Bio-Rad, Cat. No. 1708891), then diluted 1:5 in nuclease free water. Quantitative PCR was done on a Bio-Rad CFX96 Thermocycler in 96 well layout using iTaq™ Universal SYBR® Green Supermix (Bio-Rad Cat. No. 1725124). Each sample was run in duplicate in a 96 well PCR plate with each well containing 5 µL iTaq SYBR Green Supermix, 2.5 µL diluted cDNA, and 3 pMol of each forward and reverse primer. The thermal cycler running conditions were as follows: 95◦C for 30 seconds followed by 39 cycles of 95◦C for 5 seconds and 60◦C for 30 seconds. Normalized Gene Expression (NGE) was calculated using the efficiency of each primer with the following formula: [efficiency target−CTtarget/efficiency reference−CT ref]. To quantify mRNA levels, NGEs were calculated using RPL19 expression as a reference. The primers used are mentioned in table (1).

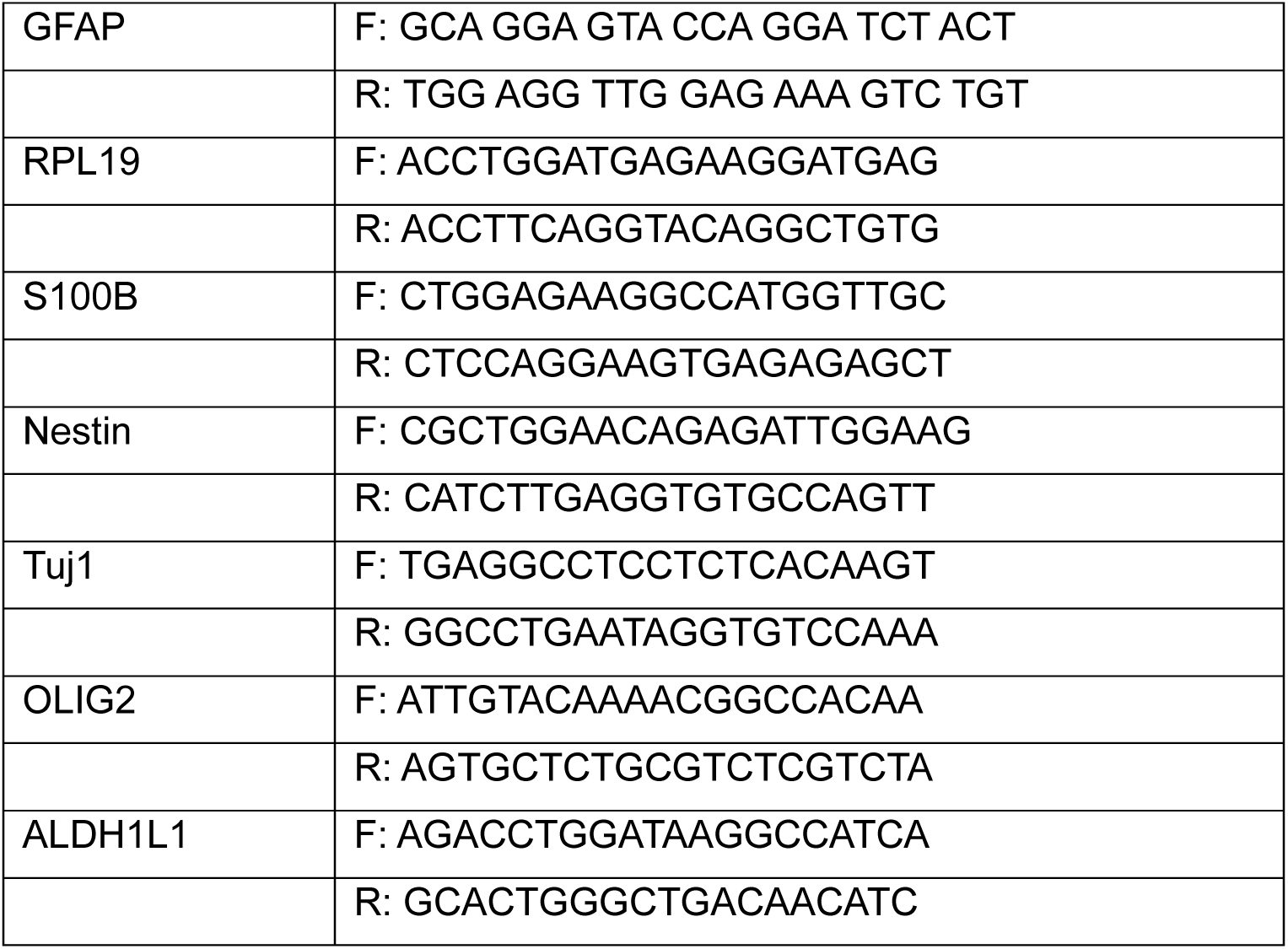

### Chromatin Immunoprecipitation -qPCR

For ChIP-qPCR, the SimpleChIP plus kit (CST, #38191) has been utilized and experiment performed according to the manufacturer’s protocols with slight modifications. Briefly, the neural precursor cells from E14.5 embryonic mouse brain of ErbB4 wild type, ErbB4 knockout, PARP1 wild type and PARP1 knockout with and without Neuregulin 1 (NRG1) treatment cultured per 10 cm dish and fixed by using 1% final concentration of formaldehyde (Sigma, #F8775) for 10 minutes at room temperature followed by 5 minutes quenching with 1X glycine (CST kit, #7005) on shaker. The fixed cells were washed with 1X cold PBS first and then scrapped after adding 1X cold PBS supplemented with 1X Protease Inhibitor cocktail (PIC); (CST, #7012). The cells then passed through 27 gauze 1ml syringe to prepare single cell suspension followed by centrifugation at 2000 g for 5 minutes at 4°C. At this step, the cells were counted through hemocytometer and equal cells (∼7 million) processed for nuclei preparation and chromatin fragmentation. The cells were incubated in 1X buffer A (CST, #7006) supplemented with1M Dithiothreitol (DTT, CST #7016) and 1X PIC for 10 minutes on rotary shaker at 4°C. Nuclei pelleted by centrifugation at 2000 g for 5 minutes at 4 °C. The supernatant removed and pellet resuspended in 1X buffer B (CST, #7007) supplemented with 1M DTT. The final nuclei pellet acquired after centrifugation at 2000 g for 5 minutes at 4 °C and processed for probe sonication. The nuclei resuspended in the lysis buffer (50 mM HEPES-KOH pH 7.5, 140mM NaCl, 1mM EDTA pH 8, 1% Triton-X 100, 0.1 % sodium deoxycholate, 0.1% sodium dodecyl sulphate, 1X PIC) and sonicated by using low amplitude of 20%, 20 sec pulse on/off cycles for 40 minutes of sonication with 1 hour interval after 20 minutes on ice (Branson SFX150, Emerson). The sonicated nuclei samples centrifuged at 10000 g for 10 minutes at 4 °C and supernatant collected as chromatin. The fragmentation analysis performed at this stage after performing reversing of cross-linking 50 ul of chromatin by using 100 ul of 1X elution buffer (CST, #7009) supplemented with 3 ul of RNase A (CST, #7013), 6 ul of 5M NaCl (CST, #7010) and 3 ul of proteinase K (CST, #10012) and incubating samples overnight at 65 °C. The samples purified through QIAquick PCR purification kit (Qiagen, #28106) and the desired chromatin fragments (100 bp to 1000 kb) were analyzed through agarose gel electrophoresis and quantified using qubit 4 fluorometer (Invitrogen). 10ug chromatin per IP sample prepared in the 1X ChIP buffer (CST, #7008) supplemented with 1X PIC and incubated with 30ul of magnetic beads to perform preclearing for 35 minutes. After that, 10% input was collected from each precleared sample and stored at −80 °C. The precleared chromatin samples were further incubated with 4ug of ErbB4 antibody (111B2; CST, #72663) and 4ug of IgG antibody (CST, #2729) for overnight at 4 °C followed by 2 hours interaction with magnetic beads at 4 °C on rotary shaker. The magnetic beads with immunocomplex washed with 1X low salt buffer (CST, # 7008) by incubating them at 4 °C on rotary shaker for 30 minutes and washing step repeated for five times. After that, 1 X high salt buffer (CST, #7008 supplemented with 70ul of 5M NaCl) wash performed for 10 minutes at 4 °C on rotary shaker. The magnetic beads separated using magnetic separator (Invitrogen DynaMag, #12321D) during each washing step and after washing. The chromatin immunoprecipitated complex exposed to the 1X elution buffer (CST, #7009) with incubation of 50 minutes at 65 °C on thermomixer F1.5 (Eppendorf, #EP5384000012). The eluent separated from the magnetic beads by utilizing magnetic separator and processed for reversing of crosslinking and purification as mentioned in the above section of chromatin fragmentation and analysis. At this stage, the purified chromatin immunoprecipitated DNA was ready for the qPCR analysis. In the qPCR, the 2ul of ChIP DNA utilized with 5ul of 1X SyBR green and reaction run for 39 cycles by keeping parameters as: Initial denaturation – 95°C for 3 minutes, Denaturation - 95°C for 30 seconds, Annealing – according to primer annealing temperature (57 – 59°C) for 45 seconds, Extension – 72°C for 30 seconds and final extension followed by melting (BioRad, CFX Opus Real-Time PCR). The designed primer list for GFAP promoter provided in the supporting information file. Results were analyzed by using CFX Maestro software and calculation of percentage input made by following equation: %Input = 100*2^[Ct Input – log2 (dilution factor) – Ct (ChIP sample)]. IgG samples were used as negative technical control, and ErbB4 knockout samples as experimental negative control to understand antibody background noise against ChIP signal. Results were represented with knockout controls.

## Statistical analysis

All statistical analyses were performed using GraphPad Prism 10.4.1. For data with two groups, paired or unpaired t-tests were used. For data with multiple groups, an ordinary one-way ANOVA with Tukey’s multiple comparisons was used for parametric statistical analysis. Kruskall Wallis followed by Dunn’s multiple comparisons to control ErbB4+/+ data, was used for non-parametric statistical analysis. For time-course PAR immunostaining, repeated measures two-way ANOVA followed by Tukey’s test was utilized. Bars for all graphs represent mean ± SEM.

## Acknowledgements and funding

We thank Dr. Judith Steen (Boston Children’s Hospital, Harvard Medical School) for assistance with proteomics; Dr. Mien-Chie Hung (The University of Texas MD Anderson Cancer Center; Graduate Institute of Cancer Biology and Center for Molecular Medicine, China Medical University, Taichung, Taiwan) for providing the phospho-PARP1 Y907 antibody; Cell signaling technologies for help in working out the chromatin IP with the ErbB4 antibody. This work was supported in part by R21NS108783, R01NS035884, P50HD105351

## Data Availability

All data generated or analyzed during this study are included in the manuscript. Original blots and microscopy images will be uploaded to Deep Blue Data, an open-access digital repository managed by the University of Michigan Library.

## Ethics Approval Statement

All animal procedures were reviewed and approved by the University of Michigan Institutional Animal Care and Use Committee; protocol PRO00012951.

